# Transgenerational dynamics of gut microbiota in black soldier fly larvae (*Hermetia illucens*) reared on a novel substrate

**DOI:** 10.1101/2025.05.28.656550

**Authors:** Shaktheeshwari Silvaraju, Amber Lim Ching Han, Tang Yong Jen, Sandra Kittelmann, Nalini Puniamoorthy

**Affiliations:** Department of Biological Sciences, National University of Singapore, 16 Science Drive 4, Singapore, 117558, Singapore; Wilmar International Limited, 28 Biopolis Road, Singapore 138568

## Abstract

Understanding the transgenerational dynamics of gut microbiota in black soldier fly larvae (BSFL) is essential for optimizing their performance on novel waste substrates in industrial settings. In this study, a wild-type BSF population was divided into six sub-lines and reared over four generations: one on standard chicken feed (CF), five on a novel diet (WIL), four of which were additionally subjected to directional selection for larval size. Despite their shared genetic origin, sub-lines exhibited divergent trajectories in larval weight and gut bacterial composition. Larval weight increased up to the second (G2) or third generation (G3) but declined sharply at generation four (G4) across all lines. Parent-offspring regressions indicated low narrow-sense heritability and minimal genetic contribution to larval weight. Gut microbiota analysis revealed that early developmental stages were most sensitive to generational shifts, with G3 to G4 transitions showing the strongest shifts in microbial communities. Notably, certain taxa such as *Bacillus* and *Paenibacillus*, involved in cellulose degradation, peaked in G2 to G3 but declined at G4, whereas *Klebsiella*, associated with immune modulation, became more abundant. These trends suggested a shift from growth-associated to digestion-oriented microbial strategies under prolonged dietary stress. However, the absence of universally beneficial taxa and the stochastic emergence of distinct microbial patterns across sub-lines highlighted the plastic and lineage-specific nature of the BSFL gut microbiota. This study emphasizes the critical need for maintaining large, genetically and microbially diverse populations in BSF breeding programs to support long-term stability and avoid performance decline when adapting to novel or suboptimal substrates.

**Importance:** The black soldier fly is increasingly used worldwide to convert organic waste into high-value protein, but the long-term stability of its gut microbiota on novel diets remains poorly understood. This study examined transgenerational changes in larval gut microbial communities from a single genetic population reared on a novel diet, with and without selection for larval size. Despite a shared genetic origin, sub-lines developed distinct microbiota and growth patterns, with early developmental stages showing the greatest sensitivity to generational microbial shifts. Initial increases in certain bacterial groups were followed by community restructuring by the fourth generation, indicating a dynamic but unstable microbial response to prolonged dietary stress. These findings highlight the importance of preserving microbial and genetic diversity when breeding black soldier flies for industrial use. Understanding how microbiota respond to selection and diet across generations is essential for sustaining performance and ensuring resilience in large-scale black soldier fly production systems.

## Introduction

The global rise in food loss and agricultural by-product accumulation presents urgent environmental and economic challenges. An estimated 1.5 billion tonnes of food waste are generated annually, contributing to 8–10% of global greenhouse gas emissions and resulting in nearly USD 1 trillion in economic losses (1). Alongside food waste, the processing of agricultural commodities generates vast quantities of underutilized waste side-streams, such as palm kernel meal (over 7 million tonnes annually; (2)) and okara, the fibrous residue of soy milk and tofu production, estimated at 14 million tonnes per year (3), just to name a few.

Insect-based bioconversion, particularly using the black soldier fly (*Hermetia illucens*, BSF), offers a sustainable solution for valorising such waste streams. Black soldier fly larvae (BSFL) efficiently convert diverse organic substrates into high-value outputs including protein, lipids, and frass, positioning them as key agents in the circular bioeconomy (4, 5). As a result, commercial interest in BSFL has expanded rapidly, with industrial-scale rearing facilities established globally (6, 7).

Optimizing BSFL production pipelines requires a deep understanding of biological processes across different stages of waste bioconversion. At the pre-valorisation stage, studies have demonstrated the importance of genetic background and diversity in shaping larval adaptability and performance (8–11). At the valorisation stage, both host genetics and diet composition have been shown to modulate larval growth, nutritional profiles, and gut microbiota composition (12–15). However, most investigations are limited to a single generation, offering only a static snapshot of host–microbe interactions in response to specific diets.

Emerging evidence suggests that BSFL can rapidly adapt to novel or suboptimal diets across generations. Transgenerational effects have been linked to improvements in larval biomass, feed conversion efficiency, and oviposition performance, including heavier and larger eggs laid over time (16–18). While these changes are often considered genetically driven, interacting factors such as diet composition (e.g., micronutrients, porosity, particle size) and the gut microbiota may play synergistic roles in shaping adaptive phenotypes (19, 20). Notably, energy allocation trade-offs between growth and maintenance, previously demonstrated in *Musca domestica*, and gene expression shifts observed in BSFL under selective pressure suggest that adaptation may involve physiological compromises; however, how such trade-offs influence gut microbial dynamics over generations remains largely unexplored (21–23).

Understanding how the gut microbiota responds to prolonged exposure to low-cost, novel substrates is critical for maintaining performance and consistency in industrial BSFL production. Although only a few studies have examined microbiota dynamics across generations, insights can be drawn from research on selective breeding, which spans multiple generations and allows exploration of microbial and host co-adaptation. For example, BSFL selectively bred for cold tolerance (16 °C) over nine generations developed a gut microbiota enriched with *Acinetobacter*, *Pseudochrobactrum*, *Enterococcus*, *Comamonas*, and *Leucobacter*. Functional metagenomics revealed upregulation of pathways related to energy and nutrient metabolism, including amino acid and lipid biosynthesis (24). Similarly, in a heat-tolerant line bred at 40 °C, the gut microbiota was dominated by *Campylobacter*, *Enterococcus*, *Dysgonomonas*, and *Proteus*, likely facilitating enhanced substrate breakdown and energy acquisition (25, 26).

While selective breeding remains a powerful tool, it is resource-intensive and may not always be practical for emerging insect production systems. the present study explored transgenerational gut microbiota dynamics in BSFL subjected to targeted, selection for increased larval size on a novel waste substrate (‘WIL’ diet comprising of 50:50 blend of palm kernel meal and okara). Using a multi-line selection experiment from a single genetic population, both phenotypic shifts in larval weight and gut microbial community composition were tracked over four generations. This framework allowed for testing whether host traits such as body weight under a novel diet drive heritable changes in the gut microbial communities. The findings aim to shed light on the transgenerational stability and adaptability of the BSFL microbiota, with implications for scalable, diet-based breeding strategies in industrial settings.

## Results

### Heritability

To evaluate the heritability of pupal weight in WT BSFL reared on a novel diet (WIL diet), average progeny pupal weights were regressed against mid-parent pupal weights across eight mating cages. Linear regression analysis yielded a narrow-sense heritability estimate of h^2^ = 0.206, based on the slope of the fitted trendline (Figure S1). The coefficient of determination was R^2^ = 0.130, indicating that approximately 13% of the variation in progeny pupal weight was explained by variation in mid-parent weight.

The heritability estimate indicated a modest but detectable additive genetic contribution to pupal weight under selection on WIL diet. Although the R^2^ value was relatively low, the presence of heritable variation suggested that the trait had the potential to respond to directional selection in this novel diet providing the basis for initiating large-scale selective breeding experiment in which the primary objective was to investigate how the gut microbiota adapted over successive generations in response to selection for larval performance on the WIL diet.

### Gut bacterial diversity influenced by diet and temporal effects over generations

The primary objective of this study was to examine the composition of the gut microbiota of a single genetic population of BSF larvae (WT) in relation to how they adapt to a non-optimal diet over four generations. A total of 9.71 million raw reads were obtained from high-throughput sequencing using ONT technology. After basecalling, demultiplexing, and quality filtering, the final dataset contained 2.81 million reads, with an average sequence yield of 15,424 ± 5,533 reads per sample (mean ± SD).

Significant differences in Shannon α diversity were observed across sub-lines (*p* = 0.0011) as well as larval age (*p* = 2.21 × 10⁻¹¹), indicating that these factors were major contributors to gut microbiota diversity. While generation alone did not significantly affect α diversity (*p* = 0.0893), significant interactions between sub-lines and larval age (*p* = 0.0014) and between generation and larval age (*p* = 0.0001) demonstrated the influence of selection and generational changes on bacterial diversity modulated over time.

To further explore these temporal trends, pairwise Wilcoxon comparisons were conducted. These revealed that Shannon diversity at experimental day 5 (D5) was consistently higher than at day 0 (D0) across multiple sub-lines. Notably, D5_WIL2, D5_WIL3, D5_WIL4, and D5_WILC all showed significantly greater bacterial diversity compared to their respective D0 counterparts (all *q* < 0.01; Figure 1). This consistent increase underscored a strong temporal effect.

**Figure 1.**
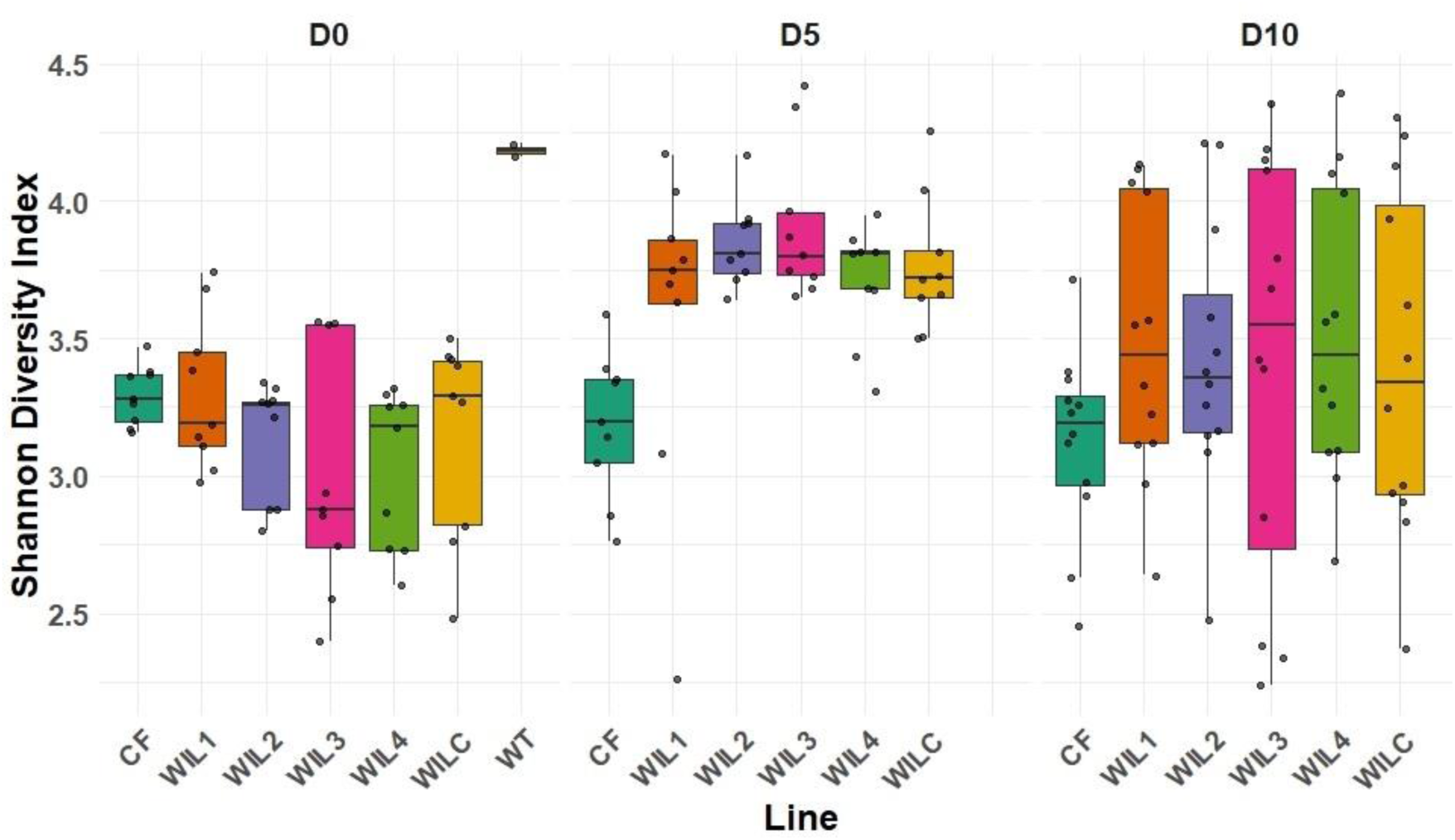
Shannon diversity index comparing bacterial communities across selectively bred sub-lines (CF, WIL1, WIL2, WIL3, WIL4, and WILC) at different experimental days (D0, D5, D10). WT D0 samples represented the initial population from which the sub-lines were derived from.

Interestingly, clear differences were observed between D0, D5, and D10_CF samples and D5 samples of sub-lines WIL2, WIL3, WIL4, and WILC (*q* < 0.05 in all comparisons), indicating a dietary influence on the bacterial diversity, particularly in BSFL fed CF diet versus those on the WIL diet. In addition, bacterial diversity in CF fed larvae remained stable across days and generations (*q* > 0.05 for all comparisons). In contrast, sub-lines undergoing artificial selection, as well as the WILC control, exhibited increased diversity at D5, further highlighting dietary pressures in shaping the gut microbiota. However, this effect was not apparent in WIL1, whereby bacterial diversity either differentiated within the line across experimental days nor between other lines (*q* > 0.05 for all comparisons).

### Rapid adaptation and diversification of gut microbiota over four generations

Principal Coordinates Analysis (PCoA) based on Bray-Curtis distances at the species level revealed distinct clustering patterns across timepoints and generations (Figure 2), which were further evaluated using pairwise PERMANOVA.

**Figure 2.**
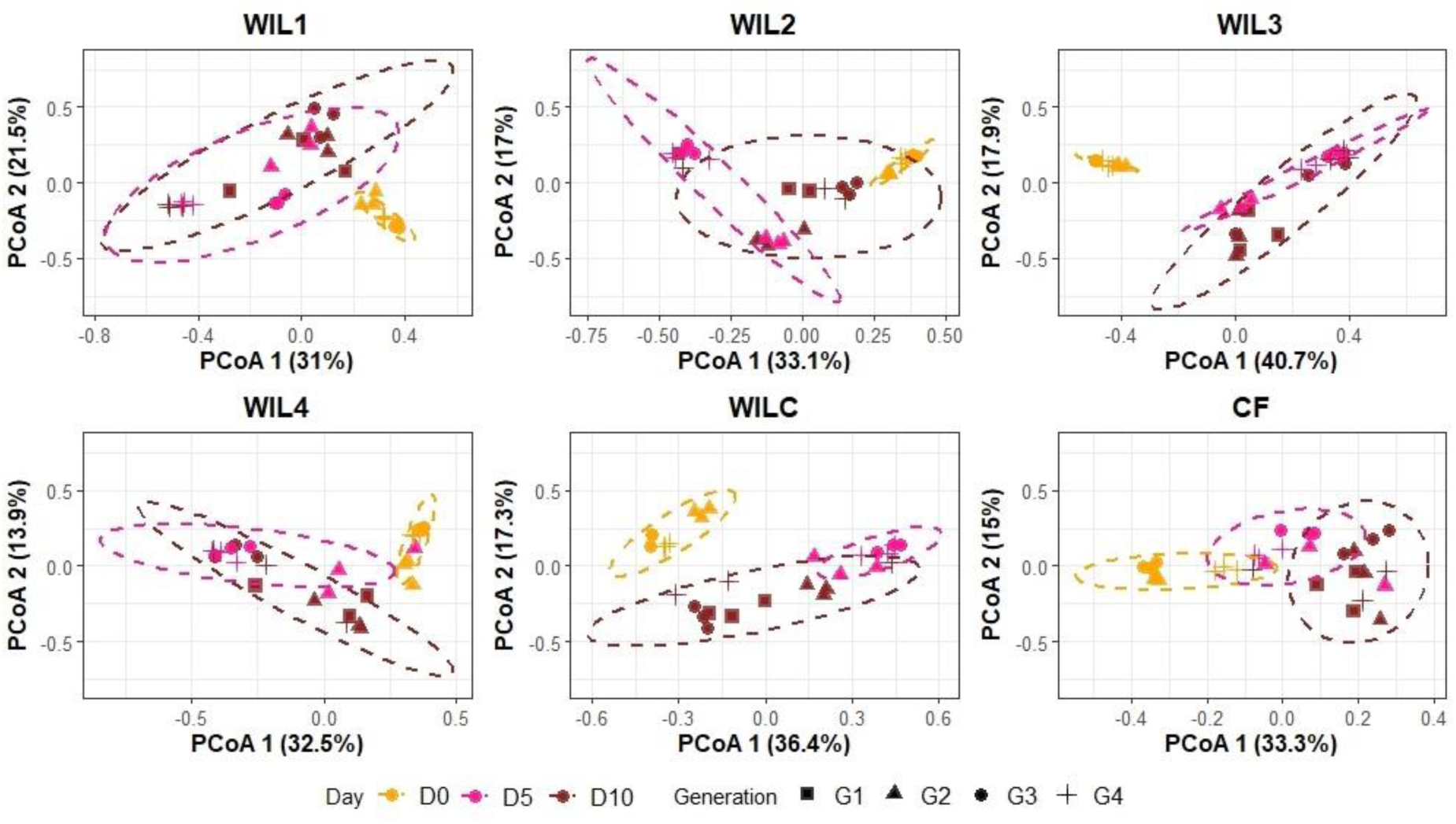
Principal Coordinate Analysis of the bacterial communities at the species level based on Bray-Curtis dissimilarity. Each panel represents a different selectively bred sub-lines (WIL1, WIL2, WIL3, WIL4) and control lines (WILC and CF). The data points are coloured according to experimental day (D0, D5, and D10) and shaped according to generation (G1 to G4). Ellipses indicate 95% confidence intervals for respective experimental day within each sub-line.

Across all populations, a strong temporal effect was evident, particularly between early and later timepoints. In every line, significant differences were observed between D0 and D5, with q-values ranging from 0.004 to 0.006, confirming that substantial microbial restructuring occurs during this early developmental window. For example, WIL1 (*q* = 0.006), WIL2 (*q* = 0.006), WIL3 (*q* = 0.006), WIL4 (*q* = 0.006), WILC (*q* = 0.004), and CF (*q* = 0.006) all showed this D0 vs D5 divergence. In most lines, significant differences were also detected between D0 and D10, again highlighting the instability of the early microbial community and its shift toward a more mature composition by later timepoints.

However, when comparing D5 to D10, the picture was more variable. Several lines, including WIL2 (*q* = 0.032), WILC (*q* = 0.004), and CF (*q* = 0.012), showed significant differences, whereas others did not. This suggests that while the initial transition from D0 to D5 is universally strong, the shift between D5 and D10 is either more subtle or plateauing, possibly reflecting the stabilization of a host-specific or diet-driven microbial community by mid- to late-adulthood.

In contrast to the consistent temporal patterns, generational effects were more selective and appeared to depend on both the line and the age. Significant generation-level differences were observed only in WIL1, WIL2, and WIL4, all of which were subjected to artificial selection. For instance, in WIL1, G2 vs G4 (*q* = 0.018) was the only generational comparison that reached significance. Similarly, in WIL2, multiple comparisons, G1 vs G2, G2 vs G3, and G2 vs G4, were significant (all *q* = 0.048), indicating a gradual yet distinguishable shift in community structure across generations. WIL4 also showed a generational effect between G2 and G3 (*q* = 0.036).

Notably, no significant generation-level differences were found in the control lines WILC and CF, reinforcing that the selective breeding process introduced microbial divergence across generations that was otherwise absent under stable, unselected conditions.

### Differential adaptation within a genetic population

To observe the taxa driving these differences in the bacterial community within the sub-lines and between the sub-lines, bacterial community structure at the phylum and genus level were investigated. Microbial community structure was profiled at three developmental time points (D0, (D5), and D10), across all sub-lines and generations (Figure 3). At D0, microbiota profiles were dominated by the phyla *Bacillota* and *Actinomycetota* across all lines. However, *Bacteroidota* was uniquely detected in the wild-type (WT) G1 sample and was absent from all other lines at this early time point, indicating distinct microbial input in the WT relative to the sub-lines. By D5, a distinct generational shift was observed in the WIL lines (WIL1 to WIL4). Notably, *Bacteroidota* emerged exclusively in G2 across all four WIL replicates, but was largely absent in G1, G3, and G4. This transient enrichment was accompanied by a concurrent increase in *Pseudomonadota* within the same G2 samples, suggesting a potential co-shift in microbial community structure specific to this generation indicating a temporary shift that was not retained in subsequent generations. At D10, microbial profiles revealed a general increase in *Pseudomonadota* and *Bacteroidota* in both CF and selected WIL lines, though their abundance remained variable. *Bacillota* showed an increase in G4 of WIL1 and WIL3 and G3 of WIL4. In contrast, *Bacteroidota* was almost absent in G2 of WIL1, WIL3, and WILC suggesting both temporal and generational shifts in community composition driven by selection and/or dietary adaptation.

**Figure 3.**
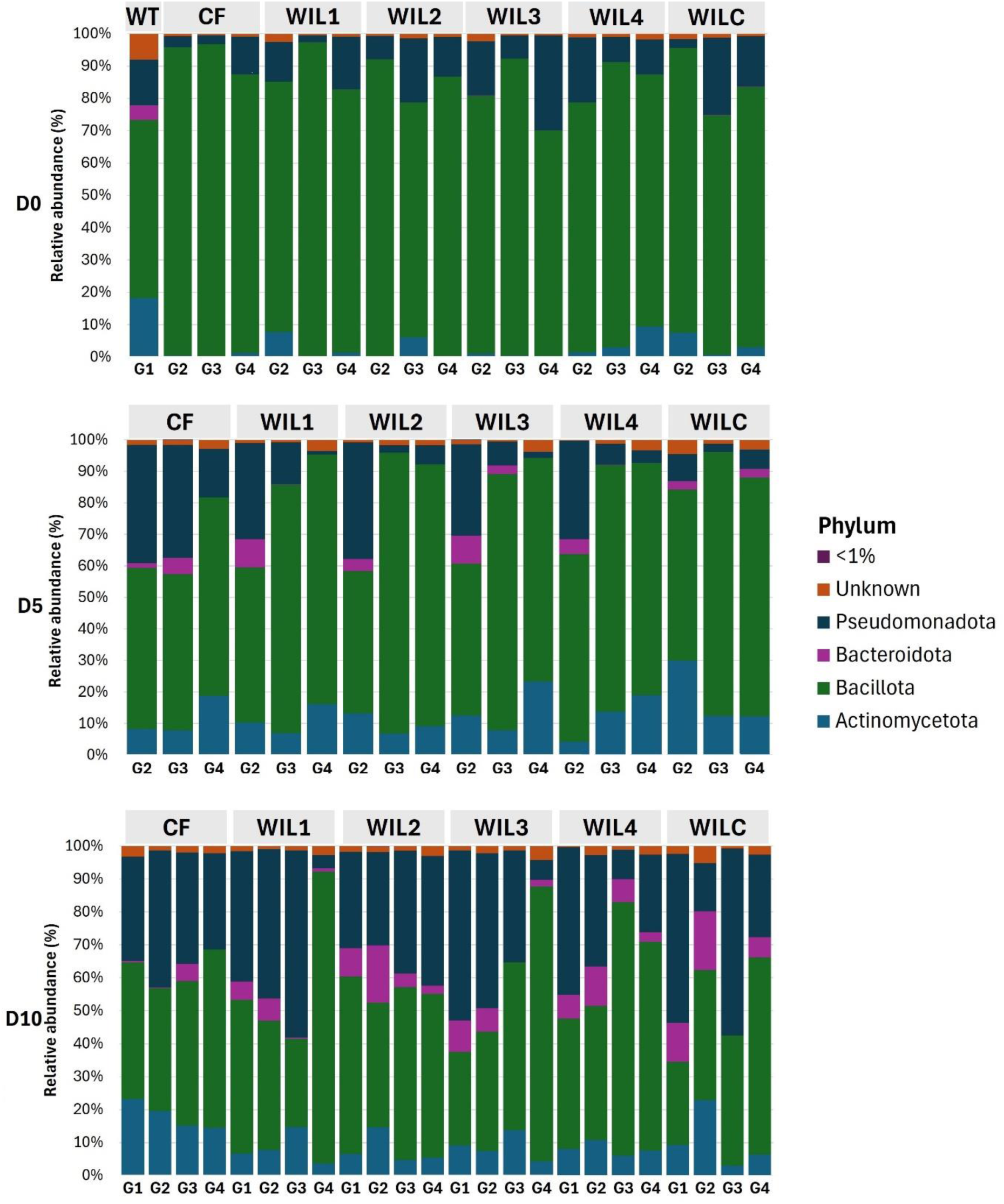
Relative abundances of phyla found in the gut of BSFL under targeted selection for weight (WIL1 to WIL4) on a novel WIL diet as well as controls (CF and WILC). “WT” represents G1, D0 samples. “<1%” includes those with < 1% relative abundance in the samples and “Unknown” includes those that was not assigned a taxon during taxonomic assignment.

To further explore microbiota differences linked to dietary selection, differential abundance analysis was performed at the genus level using ALDEx2. This revealed several taxa that significantly differed between sub-lines, with the most pronounced differences occurring between the CF control and the WIL-fed lines (WIL1 to WIL4 and WILC; Figure 4).

**Figure 4.**
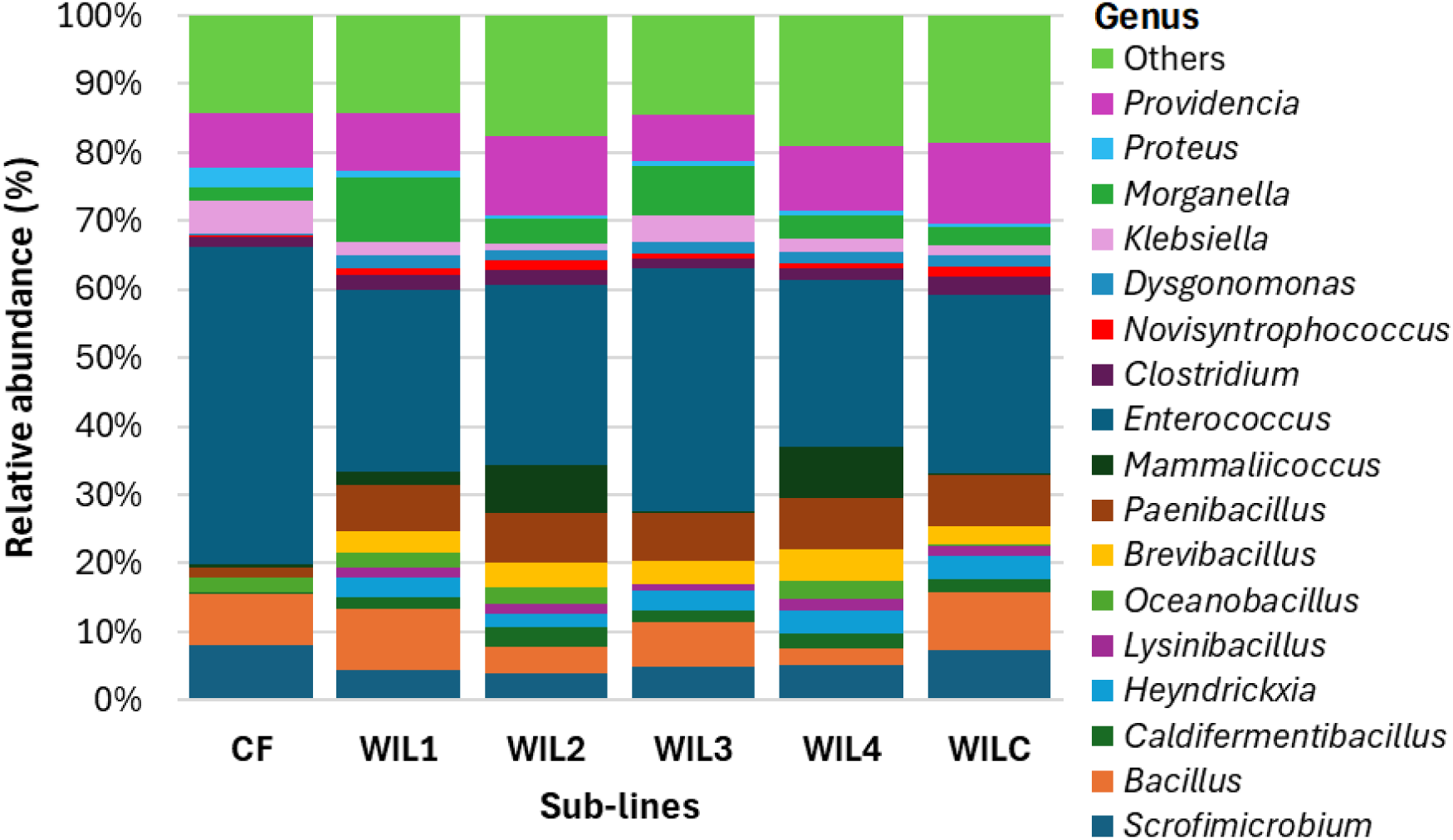
Mean relative abundances of gut-associated genera across all generations, time points, and replicates in BSFL reared under targeted selection for larval size (WIL1 to WIL4) on a novel WIL diet, alongside unselected controls (CF and WILC). “Others” represents all remaining taxa, including those with < 1% relative abundance in the samples and unassigned taxa.

Genera such as *Brevibacillus*, *Caldifermentibacillus*, *Heyndrickxia*, *Lysinibacillus*, and *Paenibacillus* were significantly enriched in all WIL-fed lines compared to CF (*q* < 0.05, Benjamini–Hochberg adjusted). In contrast, *Enterococcus* was significantly enriched in CF across all comparisons (*q* < 0.05).

Several other genera exhibited more specific patterns. *Proteus* was nearly twice as abundant in CF than in most WIL lines (2.8 ± 3.5% vs. 0.4 to 0.8 ± 0.7 to 1.6%), except WIL1, while *Klebsiella* showed higher abundance in CF compared to WIL2 (4.9 ± 5.3% vs. 0.9 ± 1.9%; *q* = 0.028). In contrast, *Novisyntrophococcus* was significantly depleted in CF relative to WILC (0.4 ± 2.1% vs. 1.4 ± 2.4%; *q* = 0.026). Notably, no significant differences in genus-level abundance were observed between any of the WIL-fed lines (*q* > 0.05).

### Temporal changes in the gut bacterial community within each generation

Although no significant genera were detected between the targeted selection lines, several taxa were differentially abundant across ages within each generation (Figure 5A) highlighting temporal changes in the gut bacterial community. Moreover, these temporal shifts were not consistent across the generations. Overall, CF displayed a stable bacterial community across ages over generations with only *Scrofimicrobium* differing between D0 and D5 in the third generation (7.4 ± 2.4% vs. 13.4 ± 8.5%; *q* = 0.048).

**Figure 5.**
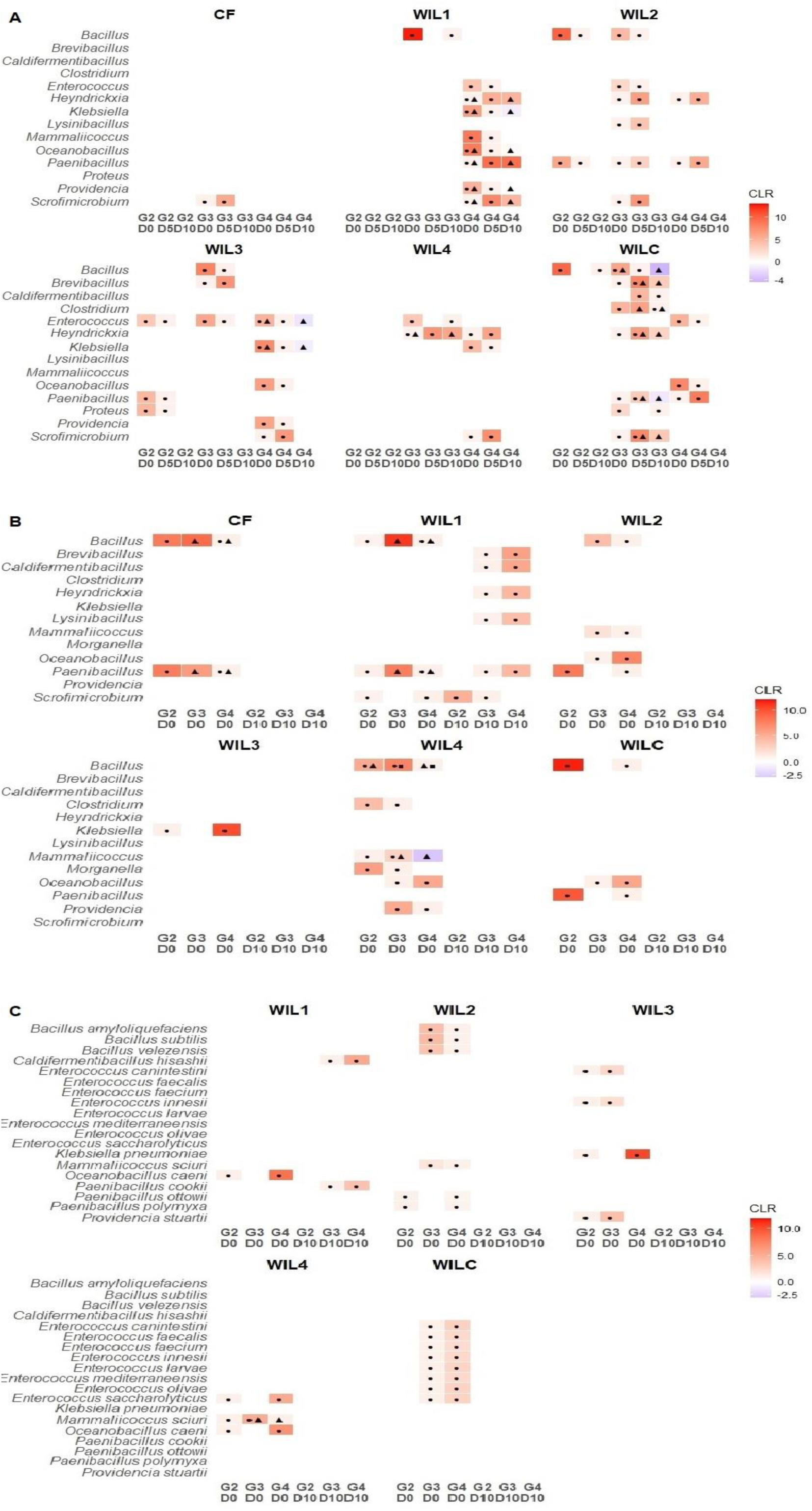
Heatmaps displaying Centered log-ratio (CLR) transformed relative abundance of bacterial taxa that were differentially abundant in the gut of BSFL across generations and/or timepoints in each sub-line. Symbol annotations (● circle, ▲ triangle, ◼ square) denote statistically significant pairwise differences (*q* < 0.05). CLR values are colour-scaled from blue (low abundance) to red (high abundance), with white representing CLR ≈ 0 (indicating q > 0.05). Only genera with a relative abundance of > 1% across all samples were included in the analysis. (A) Genera that were differentially abundant between D0, D5, and D10 within each generation (G2 to G4; G1 was omitted due to no significance detected) for each sub-line. (B) Genera that were differentially abundant between generations (G2 to G4) within each age group (D0 and D10; D5 was omitted due to no significance detected) for each sub-line. (C) Species level differences based on genera that were identified in (B); CF was omitted due to no significance detected.

Most of the temporal changes were observed in WIL1 and particularly in G4 with genera such as *Enterococcus*, *Heyndrickxia*, *Klebsiella*, *Mammaliicoccus*, *Oceanobacillus*, *Paenibacillus*, *Providencia*, and *Scrofimicrobium* differentially abundant between D0 and D5 with some of them differing between D0 and D10 as well (*q* < 0.05). The only genus that was differentially abundant in G3 was *Bacillus*, between D0 and D10 with significantly higher abundance in D10 (0.5 ± 0.4% vs., 53.6 ± 21.1%; *q* = 0.019). This genus was also observed in WIL2, WIL3 and WILC. In WIL2, temporal shifts in certain genera were observed across several generations. For instance, *Paenibacillus* differed across all generations consistently present in higher abundance in D0 compared to D5 across all the generations (*q* < 0.05). While *Enterococcus* was significantly different only in one or the other generation in WIL1 and WIL2, in WIL3, it was differentially abundant in all generations between D0 and D5 (*q* < 0.05) individuals as well as between D0 and D10 individuals in G4 (12.3 ± 6.7% vs., 66.1 ± 23.3%; *q* < 0.05). In addition, several genera including *Proteus* were significantly different within their respective generations but exclusively different between D0 and D5 (*q* < 0.05) individuals only, indicating that the bacterial community structure in WIL2 and WIL3 BSFL was relatively stable from D5 onwards with most of the bacterial shifts occurring in the early stages (between D0 and D5).

Amongst all the sub-lines that were under targeted selection, WIL4 displayed the most stability in the gut bacterial community with only *Enterococcus* significantly lower in D0 compared to D10 individuals in G3 (12.8 ± 4.3% vs., 26.5 ± 4.1%; *q* = 0.023), and *Klebsiella* and *Scrofimicrobium* differed significantly between D0 and D5 individuals in G4 (*q* < 0.05). Notably, *Heyndrickxia,* a genus that was significant in all WIL diet fed lines (except for WIL3) appeared to be differently abundant in between D0 and D5 in both G3 and G4 as well as between D0 and D10 in G3 (*q* < 0.05).

In WILC, which was not subjected to targeted selection but maintained under identical conditions, several genera not observed in other WIL sub-lines were found to be differentially abundant, most notably within the third generation. For example, *Caldifermentibacillus* and *Clostridium* were uniquely detected in WILC G3, with both genera showing significant differences between D5 and D10 (*q* < 0.05), and *Clostridium* additionally differing between D0 and D10 (*q* = 0.037), suggesting that the absence of targeted selection could have allowed relaxed selective pressures under the novel diet, facilitating shifts in gut bacterial community structure.

### The shift of specific taxa across generations

In addition to the temporal and generational influences on gut bacterial community shifts, further examination was conducted to assess how specific genera fluctuated across generations within each age group (Figure 5B), providing potential insights into generation-specific adaptation at different developmental stages.

Most of the observed differences across generations occurred at D0, suggesting that gut bacterial restructuring in response to diet was initiated early during larval development. In all the sub-lines except for WIL3 D0 individuals, *Bacillus* was observed to be differentially abundant (*q* < 0.05), however with variations in their relative abundance across generations respective to the lines. For instance, *Bacillus* was higher in abundance in G3 individuals within CF, WIL1, WIL2, and WIL4 while lower in abundance compared G4 individuals in WILC (*q* < 0.05). Similarly, *Paenibacillus* was also observed to be differentially expressed in most of the lines (except for WIL3 and WIL4) with higher abundance in G2 CF, WIL2, and WILC lines, and G3 individuals in WIL1 (*q* < 0.05). Notably, the only taxa in WIL3 that was differentially abundant between generations G2 and G4 was *Klebsiella* (2.5 ± 1.8% vs. 0.6 ± 0.4%; *q* = 0.022), and *Paenibacillus* in D5 between G2 and G3 (1.5 ± 0.9% vs. 12.9 ± 11.2%; *q* = 0.037), being the only line that showed differential abundance in any taxa between D5 and D10.

Although WIL4 exhibited minimal temporal fluctuations in gut microbial composition within each generation (Figure 5A) suggesting a relatively stable microbiota during larval development, it showed the highest number of differentially abundant taxa across generations specifically at the D0 potentially driven by primarily generational shifts in early colonisation of the gut. Specifically, *Morganella* and *Providencia*, which were not differentially present in any other sub-lines, were significantly higher in WIL4 G2 individuals compared to G3 (12.4 ± 20.3% vs. 1.4 ± 1.8%; *q* = 0.044) and in G3 individuals compared to G4 (1.8 ± 0.8% vs. 0.8 ± 0.4%; *q* = 0.026) respectively.

### Correlation between larval weight and gut microbiota

In the selection lines, larval weight generally peaked at intermediate generations before declining in G4, with the highest weights observed at G2 or G3 depending on the line (Figure S2). For instance, WIL1 larvae in G2 were significantly heavier than those in G1 (*p* = 0.0087), and both G2 and G3 were heavier than G4 (*p* = 0.0008 and *p* = 0.0002, respectively), suggesting a notable decline in the final generation. A similar trend was seen in WIL2, where G1 larvae were significantly heavier than those in G3 and G4 (*p* = 0.0005 and *p* = 0.0036), while G4 was lighter than G2 (*p* = 0.0012). In WIL3, both G1 and G2 outperformed G4 (*p* = 0.0012 and *p* = 0.0008), and in WIL4, peak weight was reached in G2, which was significantly higher than G3 and G4 (*p* = 0.0072 and *p* = 0.0008). Notably, in WILC, G3 larvae were heavier than G1 and G4 (*p* = 0.003 and *p* = 0.0007), with G2 being significantly lighter than G3 (*p* = 0.0004). These line-specific trends pointed toward a selection-driven improvement in early generations followed by possible stagnation or decline, potentially due to inbreeding depression.

To investigate the potential role of the gut microbiota in shaping larval weight, Spearman’s rank correlation was performed to identify genera associated with larval weight across generations within each sub-line. Although generation-specific testing at D10 would have provided greater resolution, the limited number of biological replicates per sub-group constrained statistical power. As a result, correlations were assessed across all generations at D10 within each line to balance interpretability and robustness. Although several genera exhibited significant correlations with larval weight (Table S2), when the D10 abundance of these correlating genera was examined across generations (Figure 5B), no consistent or significant patterns emerged except in WIL1, where multiple taxa showed differential abundance. Genera such as *Brevibacillus*, *Caldifermentibacillus*, *Heyndrickxia*, *Lysinibacillus*, and *Paenibacillus* were significantly more abundant in G4 than in G3 (*q* < 0.05), while *Scrofimicrobium* showed higher abundance in G2 compared to G3 (*q* = 0.045). Notably, several of these genera such as *Brevibacillus*, *Caldifermentibacillus*, *Heyndrickxia*, and *Paenibacillus* were among those negatively correlated with larval weight in WIL1, corresponding to the observed decline in performance in G4. However, the absence of similarly significant taxa in other lines limits the generalizability of this association, hindering conclusive interpretation of a microbiota-driven mechanism behind larval weight variation.

### Species level shifts in BSFL gut microbiota across generations

To refine genus-level patterns observed across generations (Figure 5B), species-level differences were examined within each line at D0 and D10 (Figure 5C). Consistent with genus-level results, most differentially abundant taxa emerged at D0, particularly within the lines subjected to targeted selection. These shifts were often line-specific and not conserved across developmental stages, with WIL1 being the only line to display any significant differentiation at D10. Identical to genus level observations (Figure 5B) whereby *Caldifermentibacillus* and *Paenibacillus* were differentially abundant in WIL1 D10 G3 and G4 individuals, at the species level, *Caldifermentibacillus hissashi* and *Paenibacillus cookii* were also differentially enriched between G3 and G4 individuals (*q* < 0.05), highlighting the potential role of these species in selection-driven shifts at late developmental stages over later generations.

Several *Bacillus* species *-Bacillus amyloliquefaciens*, *Bacillus subtilis*, and *Bacillus velezensis*, as well as *Paenibacillus ottowii* and *Providencia polymyxa* were differentially abundant in WIL2 between G3 and G4, and between G2 and G4 individuals with lower relative abundance observed in G4 (*q* < 0.05). Notably, WIL3 was the only line that exhibited the differential enrichment of *Providencia stuartii* (G2: 3.0 ± 2.5% vs. G3: 1.9 ± 1.4%; *q* = 0.042).

Similar to the relatively stable microbiota observed at genus level, WILC, which was not subjected to targeted selection, exhibited fewer shifts confined to early developmental stage (D0) only between G3 and G4. Furthermore, the differential enrichment was specific to *Enterococcus* with several species (uniquely expressed in WILC) such as *E. canintestini*, *E. faecalis*, *E. faecium*, *E. innesii*, *E. larvae*, *E. mediterraneensis*, *E. olivae*, and *E. saccharolyticus* showing generational differences (*q* < 0.05) suggesting that a novel diet could induce dietary adaptation influencing community composition at fine taxonomic resolution.

## Discussion

This study investigated the transgenerational effects of a novel diet on larval weight and gut microbiota in black soldier fly larvae (BSFL) derived from a single genetic population. Despite shared ancestry, sub-lines exposed to the same dietary treatment exhibited divergent trajectories in larval performance and gut microbial composition across four generations. These results underscored the dynamic interplay between environmental pressures and microbiota-mediated plasticity, demonstrating that significant phenotypic and microbial changes can arise independently of host genetic differentiation.

Larval weight trends followed a broadly consistent pattern across all sub-lines, increasing until G2 or G3 before declining sharply at G4. This trajectory was observed not only in the selection lines (WIL1–4) but also in the non-selected controls (WILC and CF), indicating that the decline in G4 was not solely the result of targeted selection. These phenotypic trends corresponded with the low narrow-sense heritability estimate for larval weight (h²), suggesting that genetic contributions to the trait were modest within this population. Instead, environmental or microbial factors, and possibly physiological trade-offs, may have played a more influential role. Previous studies have reported similar shifts in energy allocation from growth to maintenance functions in response to environmental stress, particularly under prolonged adaptation to novel substrates (16, 22).

### Transgenerational plasticity in larval gut microbiota

Microbial shifts were evident both temporally (between larval ages within a generation) and transgenerationally (across generations for a given larval age). Most strikingly, age-related changes in community composition were most pronounced in G3 and G4. *Scrofimicrobium*, a genus associated with nutrient extraction from complex organic substrates (27), was consistently enriched at D5 in nearly all lines during the later generations, suggesting a compensatory microbial response to dietary stress during peak larval metabolism. Several genera that may be associated with specific metabolic functions, such as *Enterococcus*, *Klebsiella*, *Bacillus*, and *Paenibacillus*, fluctuated significantly across the WIL diet fed lines (28). Moreover, *Bacillus*, a cellulose degrader, despite being relevant given the high cellulose content of the WIL diet (29), displayed an inconsistent pattern across the lines suggesting that microbiota adaptation occurred in a lineage-specific manner, despite uniform environmental exposure.

### Gut microbiota shift in early development

Transgenerational comparisons of each larval age further highlighted that D0 exhibited the most consistent changes across generations, suggesting that microbiota composition was influenced early in development. In particular, *Bacillus* and *Paenibacillus* were enriched at D0 in G2–G3 but declined in G4, while *Klebsiella* became highly abundant at G4. These trends likely reflected a shift in microbial functional roles rather than community destabilization. In addition, *Brevibacillus*, *Heyndrickxia*, and *Caldifermentibacillus*, genera associated with fermentation and breakdown of complex carbohydrates (30–32), were enriched in G4 D10 samples, particularly in WIL1. Although these taxa negatively correlated with larval weight in all of the WIL lines, their enrichment may represent an adaptive response wherein the microbiota reallocates functional investment from growth promotion to digestive efficiency under prolonged dietary stress, prioritising nutrient extraction over biomass accumulation, reflecting a potential trade-off strategy (19).

Notably, differentially abundant taxa were largely absent in D10 samples across most other sub-lines, despite the observed G4 performance decline supporting the idea that microbial influences on larval phenotype may be determined earlier in development. In G4 larvae, immune-modulatory species such as *Providencia stuartii* and *Klebsiella pneumoniae* were enriched (33–35), while beneficial growth associated taxa such as *Paenibacillus polymyxa* and *Bacillus subtilis* were notably reduced, suggesting a reorganization of microbiota function away from growth support (29, 36).

Interestingly, in WILC, the only control line maintained on the WIL diet without targeted selection, several *Enterococcus* species were enriched in G4, including *Enterococcus faecalis*, which has been associated with enhanced female fecundity, egg-hatching rate, and larval growth in BSFs (37). This enrichment may indicate a divergent adaptive pathway in the absence of selection pressure, favouring a microbial community structure that supported reproductive fitness and survival over larval biomass accumulation.

### Plasticity vs. long-term adaptation

Despite clear microbial and phenotypic shifts, no consistent microbial taxa emerged as universally beneficial across sub-lines or generations. This lack of convergence suggested that microbiota-mediated plasticity alone may be insufficient for long-term adaptation, particularly under strong dietary constraint. In addition, the consistent G4 decline across all lines, including CF and WILC, further supported the notion that plastic responses have inherent limits, and that sustained performance may require a stable gut bacterial community structure or co-adaptive dynamics between host and gut microbiota (38).

Importantly, the limited effective population size (∼2000 larvae per generation) may have constrained the evolutionary potential of both the host and gut microbiota. Insect systems such as *Drosophila* have demonstrated that small effective populations were prone to microbial drift, stochastic loss of beneficial symbionts, and reduced resilience to environmental stressors (38). Previous BSFL studies also suggested that improvements in performance traits under selection only stabilized after multiple generations (e.g., >9 generations), potentially due to the slow emergence of host–microbe co-adaptation under constrained conditions (24, 25).

Therefore, it is essential for industrial BSF breeding programs to account for both genetic and microbial factors influencing long-term transgenerational adaptation. Although BSFL exhibit remarkable dietary flexibility (16), successful adaptation to novel waste streams may depend on maintaining large, genetically and microbially diverse populations across multiple generations. Without sufficient genetic and microbial support, colonies may suffer from microbiota instability, reduced productivity, and eventual collapse. These findings carry practical implications for long-term colony sustainability and underscore the importance of integrating microbiota monitoring into industrial breeding strategies for BSF.

## Conclusion

This study demonstrates that transgenerational exposure to a novel diet can lead to divergent phenotypic and microbial trajectories, even in genetically identical populations. While initial improvements in larval performance may be driven by plasticity or microbial shifts, sustaining these gains over generations likely requires coordinated adaptation between host and microbiota. In the absence of such coordination particularly under small effective population sizes performance may decline, posing challenges for industrial BSFL production. These results underscore the need for integrative breeding programs that account not only for host traits but also for microbial dynamics and long-term evolutionary potential.

## Methods

### Heritability of BSF larval weight

Wild caught BSF were used to establish a stock population (WT), which had a low observed inbreeding coefficient (Fis = 0.121) prior to the commencement of the experiment (6). Narrow-sense (additive) heritability (h^2^) of pupal weight was estimated using a single-generation parent–offspring regression approach. Fifty 5-day-old WT larvae were randomly selected and transferred into a plastic container containing the WIL diet, composed of 50% palm kernel meal (PKM) and 50% okara (OKA) (sourced from Wilmar International Pte Ltd) with a moisture content of 70%. Upon pupation, each larva was individually weighed and isolated in separate *Drosophila* breeding plastic tubes to prevent uncontrolled mating.

After adult emergence, individuals were sexed, and each was retained in isolation for an additional four days to allow females to reach reproductive maturity. The heaviest male and female were selected and introduced into mesh mating cages (each pair per cage, total 15 pairs) containing a corrugated cardboard strip suspended above an oviposition attractant composed of chicken feed and frass. Cardboard strips with deposited eggs were removed and suspended above fresh chicken feed in a new container, allowing hatched larvae to drop directly onto the feed.

Five-day-old progeny were collected and subjected to the same treatment as the parental generation. Both parental and progeny individuals were weighed at the pupal stage. Mid-parent pupal weight was calculated as the average of the male and female parental weights for each mating pair. The mean pupal weight of the progeny from each pair was then calculated from 10 pupae derived from per mating pair and used for heritability estimation. Narrow-sense heritability was estimated using parent–offspring regression, applying the equation:

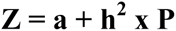

where *Z* represents the average pupal weight of the progeny, *P* is the mid-parent pupal weight, *a* is the intercept, and *h^2^* is the slope of the regression line, representing the additive genetic contribution to trait variation. The coefficient of determination (R^2^) was calculated to indicate the proportion of variance in progeny weight explained by mid-parent weight.

Linear regression was performed in R (version 4.1.2) using the lm() function, with mid-parent pupal weight as the independent variable and average progeny pupal weight as the dependent variable. The slope of the regression line (h²) represented the proportion of phenotypic variance attributable to additive genetic variance. A total of eight families were included in the analysis. **Experimental set-up**

Following the heritability estimation, a large-scale diet experiment was conducted to investigate the transgenerational response of larval size and gut microbiota to a novel substrate. The wild-type (WT) founder population was divided into six sub-lines, each with three replicates: CF, WILC, WIL1, WIL2, WIL3 andWIL4. The CF sub-line functioned as a general control and was reared on a standard chicken feed (PK Agro-industrial Products M Sdn Bhd; Johor, Malaysia) whilst WILC functioned as diet-specific control (without selection), reared on a novel diet (WIL diet: 50% palm kernel meal (PKM), 50% okara (OKA)). Sub-lines WIL1 to WIL4 were subjected to directional selection for larger larval size on the WIL diet. Each replicate of the sub-lines was initiated with 2000 WT larvae and reared on their respective diets provided *ad libitum* (6.678 kg total wet weight per tray). At approximately day 10, when more than 50% of larvae reached the prepupal stage, they were subjected to a two-step sieving process to separate larvae from residual substrate. The initial sieving was performed using an industrial-grade sieve with a mesh size of 3 mm, followed by manual sieving with a slightly smaller mesh size of 1 mm to further remove debris.

For the size selected lines (WIL1–WIL4), a third sieving step (mesh size: 5 mm) was implemented to separate larger larvae/prepupae from smaller individuals. From these, the top 1000 largest individuals were selected for adult emergence and mating to produce the next generation. The CF and WILC sub-lines experienced no size selection; instead, a subset of 1000 larvae/prepupae were randomly chosen for mating without regard to size. This was repeated over three successive generations (G2 to G4) to monitor phenotypic response, and gut microbiota shifts under diet-mediated selection pressure.

### Sample collection and gut isolation

Three WT larvae were collected at the experimental start date (D0) and snap frozen on dry ice. Ten larvae were collected from each of three replicates from each sub-line (CF, WIL1 to WIL4, and WILC) on day 5 (D5), day 10 (D10), weighed and snap frozen as well to analyse the changes in the gut microbiota over time. Most of the larvae began entering prepupal to pupal phase from D10 to D15. Sample collection was performed at every generation (first generation: G1 to fourth generation: G4) and all samples were stored at −80°C until further processing.

The dissection procedure followed the technique outlined in previous studies (15, 39). Three whole larval guts per replicate were excised and pooled together to ensure sufficient material for subsequent DNA extraction. Before and after each excision, 70% ethanol was used to sterilize all surfaces and instruments. Isolated gut samples were stored at −80 °C until further processing.

### Microbial community profiling

DNA was extracted from isolated gut samples using the QIAamp^®^ PowerFecal^®^ Pro DNA Kit (QIAGEN) following the manufacturer’s handbook, employing the vortex adapter method for the homogenization of tissue samples. Before the final centrifugation step, 50 µl of C6 solution were added onto the MB Spin Column filter membrane and the column left to incubate at room temperature for 5 min to ensure sufficient time was provided for the DNA to completely dislodge.

Polymerase chain reaction (PCR) amplification was done targeting the full length 16S rRNA gene (V1-V9 regions) using custom designed barcoded 27F and 1492R primer pairs (Table S1) with unique barcode combinations for multiplexing. 10 µl of Q5^®^ High-Fidelity DNA Polymerase (New England Biolabs, Singapore), 1 µl of forward and reverse pre-mixed primer pairs with unique barcode combinations each (10 µM), 50 to 100 ng DNA template, and water to reach a final volume of 25 µl were used. Each reaction was set up in triplicates to minimise amplification bias and no template controls were included to ensure absence of contamination. The PCR cycling conditions were as follows: denaturation for 2 min at 95°C, 30 cycles of 20 s at 95°C, 20 s at 55°C, 1 min at 72°C, followed by final extension at 72°C for 7 min (Bio-Rad T100 Thermal Cycler). Verification of successful amplification was conducted through gel electrophoresis. PCR amplicons (∼1500 bp) were quantified and pooled in equimolar concentrations to generate seven libraries, each comprising 26 samples. These pooled libraries were subsequently prepared and barcoded using the Native Barcoding Kit 24 V14 (Oxford Nanopore Technologies), following the manufacturer’s instructions. Final libraries were loaded onto MinION Flow Cells (R10.4.1, FLO-MIN114) and sequenced using the Oxford Nanopore Technologies MinION platform.

### Nanopore bioinformatics

Following sequencing on the Oxford Nanopore platform, raw reads were basecalled and were first demultiplexed using Guppy to separate libraries based on Native Barcodes (NBD). Subsequently, demultiplexed reads from each library were subjected to a second round of demultiplexing and barcode trimming using a custom MATLAB script (Supplementary file: demul_and_trim_BothStrands.m) based on barcoded primers appended to individual samples. Sequencing reads in FASTQ format were imported and scanned for sample-specific barcodes provided in an accompanying barcode mapping file (Supplementary file: 16s_barcode sequence for matlab.xlsx).

Demultiplexing was performed by detecting matching barcode sequences within the first and last 100 base pairs of each read. Reads were assigned to a specific sample only if both forward and reverse barcodes were identified. Once assigned, barcode regions were trimmed. Additional filtering was applied to retain reads within the size range of 1,000 to 2,000 base pairs to ensure high-quality, near-full-length 16S rRNA sequences.

Basecalled and demultiplexed reads exported as individual FASTQ files per sample were uploaded to the EPI2ME platform (version 2.11.0, Oxford Nanopore Technologies) for taxonomic classification using the Metagenomics Workflow. Within this pipeline, reads were aligned against the reference database using Minimap2 (version 2.26-r1175) as the classification engine. Taxonomic assignments were resolved down to the species level. The pipeline generated raw read count tables and diversity metrics, including Shannon diversity index values, which were exported for downstream statistical analysis.

### Statistical analysis

#### Statistical analysis of larval weight

Larval weight on experimental day 10 (D10) were analysed to assess generational differences within each sub-line. Statistical analysis was performed using R (version 4.1.2). For each line, a one-way Analysis of Variance (ANOVA) was conducted to test for significant differences in larval weight among generations. When a significant overall effect was detected (α = 0.05), Tukey’s Honest Significant Difference (HSD) test was applied as a post hoc analysis to evaluate pairwise differences between generations. This method accounts for multiple comparisons by controlling the family-wise error rate, and the resulting adjusted *p*-values (*q* values) were used to assess significant differences in larval weight between generations within each sub-line followed by visualization using box-and-whisker plots using ggplot2 (version 3.5.1).

#### Statistical analysis of 16S rRNA gene sequence data

Raw count abundance tables, taxonomic assignments, and α diversity metrics (Shannon index) generated via the Epi2Me platform were imported into RStudio (version 4.1.2) for downstream statistical analysis. All analyses were performed using a combination of R packages, including phyloseq, vegan, ggplot2, dplyr, tidyverse, patchwork, and ALDEx2.

Shannon α diversity values were analysed using pairwise Wilcoxon rank-sum tests between groups (sublines, generations and larval age), with *p*-values adjusted using the Benjamini– Hochberg method (reported as *q* values). A three-way ANOVA was additionally performed to assess the main and interaction effects of sub-lines, generations and larval age on Shannon α diversity. Boxplots were generated to visualize diversity changes over time and across experimental groups.

To evaluate differences in overall community structure within each sub-line, β diversity was assessed using Principal Coordinates Analysis (PCoA) based on the Bray–Curtis dissimilarity metric, clustering with 95% confidence ellipses grouped by age (D0, D5, D10) and generation (G1–G4) (phyloseq package version 1.38.0).

Permutational Multivariate Analysis of Variance (PERMANOVA) was performed using the vegan package (version 2.6-8), with 999 permutations to test for the effects of sub-lines, larval age, and generations on microbial community composition. PERMANOVA was applied in three contexts: (1) globally across all generations and timepoints within each line, (2) pairwise comparisons of days within each generation and generations within each day, and (3) comparisons including WT samples (G1, D0) as a baseline reference. Pairwise PERMANOVA *p*-values were adjusted for multiple testing using the Benjamini–Hochberg (BH) correction method reported as *q* values.

Differential abundance analysis was conducted using the ALDEx2 package (version 1.26.0). Abundance data (with taxa < 1% across all samples collapsed) were centered log-ratio (CLR) transformed and underwent Monte Carlo Dirichlet sampling with 128 Monte Carlo instances (mc.samples = 128). Statistical testing was performed using Welch’s t-test, and significance was determined based on the Benjamini–Hochberg-adjusted *p*-values (*q* < 0.05). Pairwise comparisons were conducted (1) between sub-lines, (2) across timepoints within each generation, and (3) across generations within each timepoint at both genus and species level. Heatmaps were generated using ggplot2 (version 3.5.1) to visualize CLR-transformed values for taxa meeting the abundance threshold, with symbol annotations indicating statistically significant pairwise differences.

Spearman’s rank correlation coefficients were calculated using Microsoft Excel to examine the relationship between larval weight and gut bacterial taxa at both genus and species levels across sub-lines. Statistical significance was assessed using Student’s *t*-test, with adjustments for tied ranks applied using the Bonferroni correction method.

## Acknowledgements

This research was jointly funded by the National Research Foundation of Singapore (NRF2020-THE003-0003/A00085140000), the Ministry of Education Singapore (A00044240000), and the Economic Development Board – Industrial Postgraduate Programme (EDB-IPP) jointly awarded to Wilmar International Limited and the National University of Singapore (A00085100000). The funders had no role in study design, data collection and interpretation, or the decision to submit the work for publication.

